# Analytical resolution governs cross-laboratory and cross-species concordance in murine cardiometabolic HFpEF transcriptomics

**DOI:** 10.64898/2026.04.30.721824

**Authors:** Amin Forouzandehmehr

## Abstract

**Background.:** Murine cardiometabolic heart failure with preserved ejection fraction (HFpEF) models are selected and generalized from pathway-level transcriptomic summaries, but how reproducibly those signals hold across laboratories and species is unclear.

**Methods.:** I re-analyzed ventricular RNA-sequencing, ventricular single-cell data and subgroup-resolved human HFpEF proteomics for db/db plus aldosterone (db/db+Aldo) and high-fat-diet plus L-NAME (HFD+L-NAME) models, quantifying agreement at gene and pathway level with bootstrap intervals, and applied the identical pipeline to five mouse cardiac model pairs, each two independent deposits at matched sample size.

**Results.:** For HFD+L-NAME, agreement across 23,838 genes was rho = 0.08 (95% CI 0.06– 0.09), rising to 0.45 among 349 differentially expressed genes, whereas the eight-pathway summary gave rho = 0.71 with an interval spanning −0.04 to 0.98. A transverse aortic constriction positive control at matched sample size reached rho = 0.54 at gene level and 0.90 at pathway level, with 2,241 genes significant at four mice per arm. Across five model pairs gene-level agreement ranged nearly nine-fold (0.08 to 0.69) while pathway-level agreement never fell below 0.71. Cross-species concordance was confined to oxidative phosphorylation (142 genes). Independent single-cell projection placed the myofibroblast signature in mural cells rather than fibroblasts.

**Conclusions.:** Apparent reproducibility in murine HFpEF transcriptomics is governed by the resolution at which it is measured. Pathway-level correlations were uniformly high, including for models that do not reproduce gene by gene, and cannot certify reproducibility. Reporting gene-level agreement, feature support and interval estimates turns public omics into usable evidence for model selection and 3Rs-informed design.

## Introduction

Heart failure with preserved ejection fraction (HFpEF) is a major and growing form of heart failure characterized by diastolic dysfunction, myocardial remodeling, and frequent coexistence of obesity, diabetes and other cardiometabolic comorbidities [1,2]. Because human myocardial tissue is scarce, mechanistic work depends heavily on murine models. Two are among the most widely used: leptin receptor-deficient db/db mice treated with aldosterone (db/db+Aldo), and high-fat diet combined with the nitric oxide synthase inhibitor Nω-nitro-L-arginine methyl ester (HFD+L-NAME) [3,4].

Choosing between such models is consequential. A model chosen because it is believed to reproduce a particular human pathway determines which mechanisms are pursued, and how many animals are consumed pursuing them. That choice is usually justified by transcriptomic evidence summarized at the level of pathways or gene-set scores, on the reasoning that individual genes are noisy while pathway aggregates are stable.

Public datasets now make it possible to test this assumption directly. Aggregation can reduce noise and reveal coordinated biology, while its evidential strength depends on the number and dependence of the features being compared. A resolution-aware analysis can therefore distinguish robust biological signal from summary-level amplification and determine which conclusions transfer most consistently across datasets.

Here I introduce a resolution-matched benchmarking framework for published murine and human HFpEF datasets. By comparing the same mouse model across independent laboratories and two mouse models against subgroup-resolved human HFpEF proteomics at gene, pathway and cell-program levels, I define where reproducibility resides and which signals translate to human disease. The analysis identifies substantial replication among the strongest gene-level effects, calibrates the framework against a transverse aortic constriction positive control profiled at matched sample size, extends the same measurement to five mouse cardiac model pairs, resolves a cross-species oxidative-phosphorylation signal in the HFD+L-NAME model, and provides a practical single-cell validation test for bulk signatures. Together, these results turn heterogeneous public omics datasets into a ranked evidence base for HFpEF model selection.

This framework has direct translational and 3Rs value. It identifies the model features most likely to reproduce, distinguishes well-supported cross-species signals, and allows public data to guide experimental design before additional animals are committed.

## Methods

### Data sources

Murine bulk RNA-sequencing and matched echocardiographic and morphometric data were obtained from GSE284354 and its associated publication [4]. That deposit contains two cardiometabolic HFpEF preparations. The db/db+Aldo cohort comprises 15 animals (8 db/db+Aldo, 7 wild-type vehicle controls; 8 female and 7 male). The HFD+L-NAME arm comprises 8 animals (4 per group). An independent HFD+L-NAME dataset, GSE180065 [5], generated with the model as originally described [6], comprises 15 animals (5 chow controls; 10 HFpEF, five at five weeks and five at seven weeks), and was rebuilt for this study directly from the per-sample files in the GEO deposit. Ventricular interstitial single-cell RNA-sequencing data were obtained from GSE275031 [7]. Subgroup-resolved human HFpEF myocardial proteomics were obtained from the public deposit described in [8]. A positive control was analyzed with the same pipeline. Two independent transverse aortic constriction (TAC) datasets were used: GSE322779 [9], comprising left ventricular counts from wild-type littermate controls (4 sham, 4 TAC), and GSE283181 [10], comprising left ventricular expression values from wild-type mice (4 sham, 5 TAC). Both were chosen before analysis against fixed criteria: the same model in two independent laboratories, ventricular bulk RNA-sequencing, a wild-type control arm, sample size matched to the HFD+L-NAME comparison so that reproducibility could not be confounded with statistical power, and depositor-assigned group labels. Deposits screened against these criteria and not used are listed with the code. To test whether the resolution effect generalizes, the same two measurements were repeated on three further mouse cardiac models, each represented by two independent deposits with depositor-assigned labels and three to six animals per arm: doxorubicin cardiotoxicity (GSE224157 and GSE307421), isoproterenol infusion (GSE217755 and GSE255532) and myocardial infarction (GSE275835 and GSE304090). These cohorts are identified by accession throughout.

### Validation of sample group labels

Group assignments were verified against two independent sources before any contrast was computed. First, the morphometric and echocardiographic data separate the db/db+Aldo cohort completely: body weight 38.9-59.5 g versus 21.1-31.1 g, blood glucose 343-600 versus 132-198 mg/dL, and E/e’ 47.6-68.1 versus 21.4-29.5, with no overlap on any measure. Second, recomputing the group contrast from the deposited expression matrix under these labels reproduces the depositors’ own DESeq2 fold changes (Spearman rho = +0.993, 95.9% sign agreement across genes with adjusted p below 0.01). Labels for the HFD+L-NAME arm were verified the same way (rho = +0.988, 100% sign agreement). This verification runs as a mandatory first step in the released pipeline, and I recommend the practice generally: label errors are invisible to downstream statistics and are detected only by checking against an independent measurement.

### Pathway activity and cross-dataset comparison

Pathway activity was quantified by single-sample gene set enrichment analysis using the GSVA package [11,12]. Mouse Hallmark [13,14] and Gene Ontology Biological Process [15] sets were obtained from MSigDB; custom excitation-contraction coupling, ion-channel and mechanotransduction sets were curated from cardiac mechanobiology [16] and cardiac physiology [17–19] literature. All gene sets used, together with cell-type marker lists and the transcription-factor regulon table, are released as plain files with the analysis code.

For cross-laboratory comparison, both cohorts were reduced to a common gene universe before scoring, because single-sample enrichment ranks genes within a sample and is therefore not comparable across datasets built on different gene universes. Agreement was then quantified at two levels: across all shared genes, and across the eight pathway summaries. Because the two deposits are normalized differently, directional agreement is reported both raw and after median-centering the effect distributions.

For cross-species comparison, murine gene-level effects were correlated with human proteomic effects across the genes shared between a given pathway and the human deposit. Every comparison reports the number of shared genes on which it rests. A comparison was treated as interpretable only where at least 15 genes overlapped; this pragmatic threshold was fixed before results were examined. Correlations are reported with bootstrap 95% confidence intervals (2,000 feature resamples) and permutation p-values (5,000 permutations). Because genes within pathways are correlated, these intervals quantify sensitivity to the observed feature composition and should not be interpreted as additional biological replication.

### Cell-program signatures

Curated marker sets were scored by single-sample enrichment in bulk data as previously described [20–22], and the same sets were projected onto the independent single-cell dataset using module scoring in Seurat [23]. Marker-set specificity was additionally assessed directly, by tabulating the overlap between curated sets and by classifying each member of the myofibroblast set against canonical lineage markers.

### Severity modeling

Elastic-net models of a phenotype-derived remodeling severity score were fitted by elastic-net regression [24] implemented with glmnet [25] under nested leave-one-out cross-validation, and are reported in the supplementary material only. Out-of-sample performance is reported as 1 − SSE/SST, the residual sum of squares over the total sum of squares, which can be negative when a model performs worse than the sample mean; the squared observed-predicted correlation, which cannot fall below zero and therefore cannot reveal a model without skill, is reported alongside for comparison. Because the candidate pool differed by two orders of magnitude between feature classes, the comparison was repeated with the transcription-factor pool randomly subsampled to the number of pathway features. The corresponding analyses are presented in Figure S1.

### Statistics

Analyses were performed in R 4.6.1. Group comparisons used the Wilcoxon rank-sum test and correlations the Spearman coefficient. One multiple-testing rule was applied throughout: Benjamini-Hochberg correction within each family of tests reported together, with the family stated in each case. Adjusted p below 0.05 was called significant; everything else was described as a trend or nominal signal rather than an established effect. Uncertainty on correlation estimates is given as bootstrap 95% confidence intervals. For correlations across genes or pathway summaries, resampling was performed over the compared features; these intervals describe feature-level stability and do not replace replication at the animal or cohort level. Permutation p-values were obtained by randomly permuting one of the two compared effect vectors and comparing the absolute observed correlation against the resulting null distribution; 2,000 permutations were used throughout except for the cross-species comparisons, which used 5,000. These are reported as nominal, two-sided values.

## Results

### Cross-laboratory reproducibility concentrates in the largest transcriptional effects

I first mapped cross-laboratory reproducibility of the HFD+L-NAME response in two independent cohorts (GSE284354 and GSE180065). Across the complete universe of 23,838 shared genes, the transcriptome-wide baseline was Spearman rho = 0.078 (bootstrap 95% CI 0.064 to 0.092), with 52.8% raw and 54.4% median-centered directional agreement (Figure 1A, B). This unfiltered benchmark allowed the reproducible high-signal component to be identified directly.

**Figure 1.**
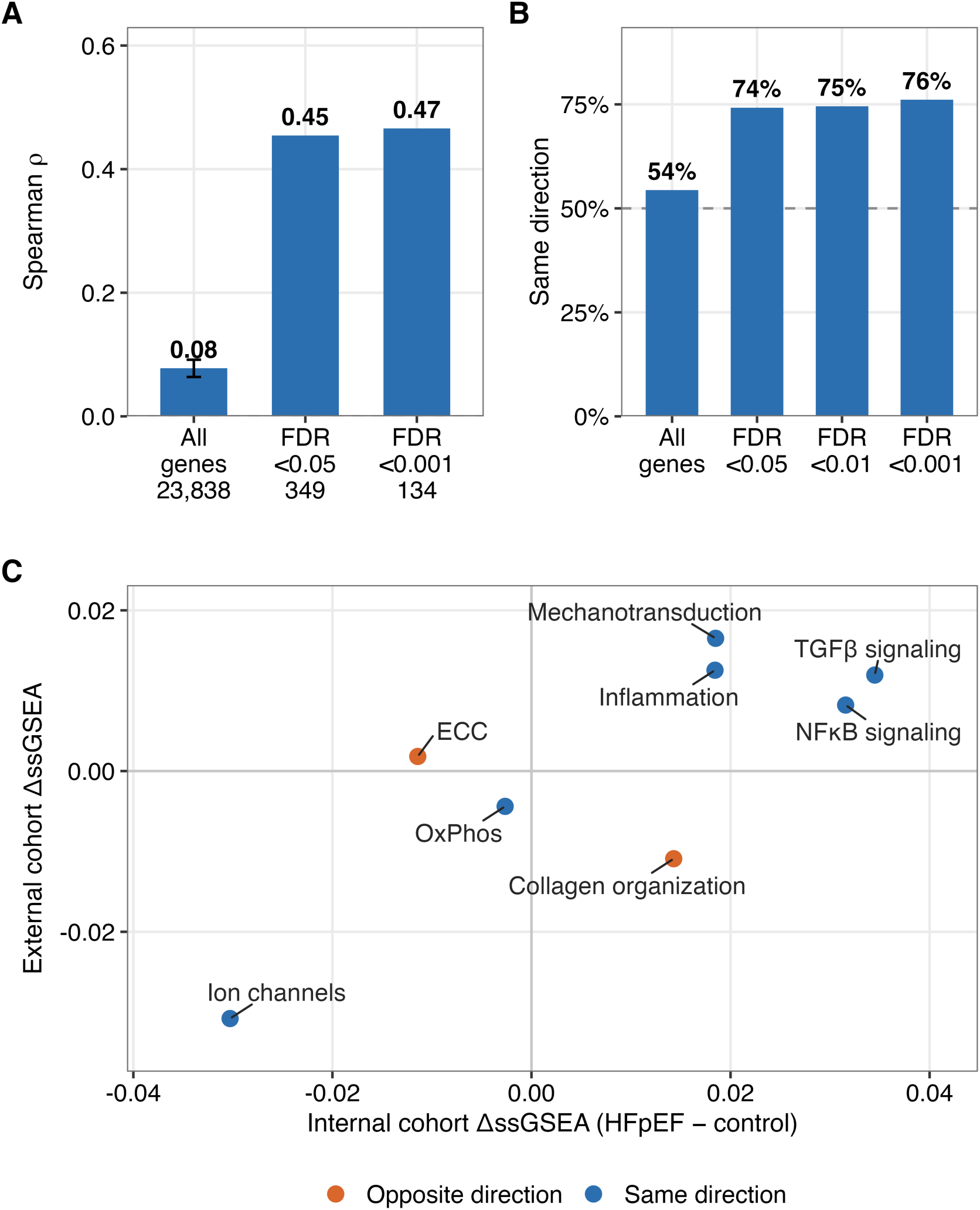
Cross-laboratory reproducibility of the HFD+L-NAME model. (A) Spearman correlation between per-gene HFpEF-versus-control effects in two independent cohorts (GSE284354 and GSE1800C5), across all shared genes and restricted to genes called differentially expressed in the source study at the thresholds shown. Error bar, bootstrap S5% confidence interval. (B) Fraction of genes changing in the same direction in both cohorts after median-centering; dashed line, chance. (C) The same comparison summarized as eight pathway activity scores. Blue, pathways agreeing in direction; orange, disagreeing.

Effect-support stratification revealed that reproducible component. Among the 349 genes called differentially expressed at adjusted p below 0.05 in the source study, independent-laboratory agreement rose sharply to rho = 0.454 with 74.2% directional concordance. Concordance remained substantial and directional agreement increased at more stringent thresholds (216 genes at adjusted p below 0.01, rho = 0.441, 74.5%; 134 genes at adjusted p below 0.001, rho = 0.466, 76.1%; Figure 1A, B). The strongest effects therefore replicated under independent laboratory conditions even in the compact four-per-group cohort.

Cross-laboratory reproducibility is therefore concentrated in the biologically strongest effects: nearly three quarters reproduced in direction, with concordance increasing as statistical evidence strengthened. This defines a robust transcriptional core of the HFD+L-NAME model.

### Pathway-level summaries yield large correlations with wide confidence intervals

Summarizing the same two cohorts as eight pathway activity scores gave Spearman rho = 0.714, p = 0.058, with six of eight pathways shifting in the same direction (Figure 1C). The estimate is large, and it rests on eight points.

That estimate is weakly constrained: its bootstrap 95% confidence interval spans -0.04 to 0.98 because it rests on eight pathway points. The same comparison across 23,838 gene-level features gives rho = 0.078 with a much tighter feature-resampling interval. Although correlated genes are not independent biological replicates, the joint view shows why effect magnitude, feature support and precision are complementary. Reporting all three preserves the biological insight of pathway concordance while making its evidential strength explicit.

Pathway-specific direction further sharpens the biological interpretation. Six programs agreed across laboratories, whereas collagen fibril organization shifted upward in the internal cohort and downward in the external one, and excitation-contraction coupling also differed in direction. The framework therefore pinpoints both conserved and context-sensitive biology rather than collapsing them into a single summary.

### A transverse aortic constriction control separates gene-level from pathway-level agreement

The weak gene-level agreement between the two HFD+L-NAME cohorts admits two readings: the model may reproduce poorly at gene level, or the framework may be unable to resolve gene-level agreement at four to five mice per arm. I separated them with a positive control, applying the identical pipeline to two independent transverse aortic constriction cohorts held at the same sample size (GSE322779 [9] and GSE283181 [10]). Across 16,538 shared genes the cross-laboratory correlation was rho = 0.535 (bootstrap 95% CI 0.522 to 0.547; permutation p < 0.001), with 65.6% of genes agreeing in direction, against rho = 0.078 for HFD+L-NAME (Figure 2A, B). Restricting to genes called differentially expressed in the reference cohort raised agreement to rho = 0.748 at adjusted p below 0.05 and rho = 0.610 at adjusted p below 0.001, with 90% and 93% directional agreement.

**Figure 2.**
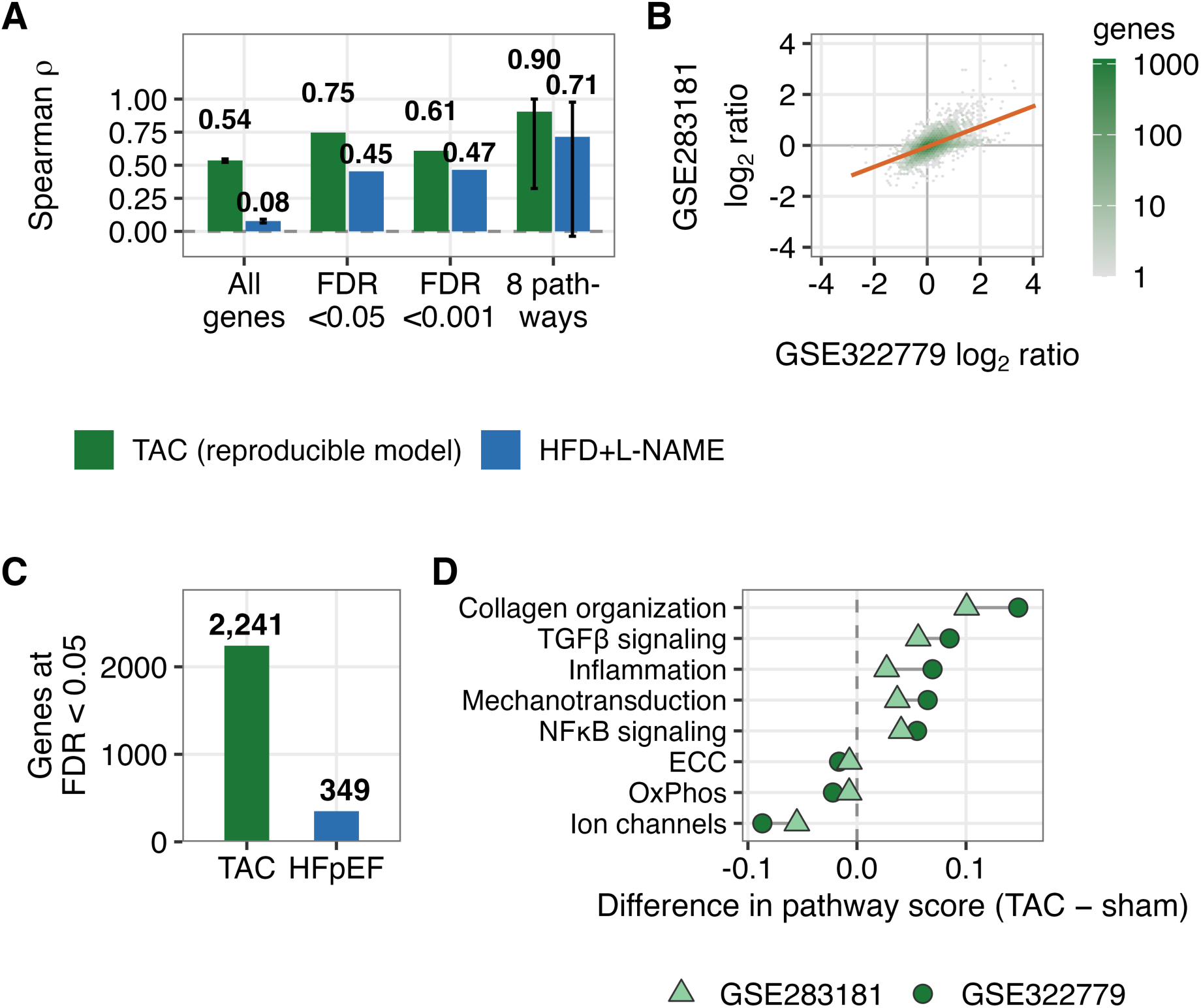
Positive control: cross-laboratory concordance for a model that reproduces. Two independent transverse aortic constriction (TAC) cohorts, GSE32277S [S] and GSE283181 [10], were analyzed with the pipeline applied to the HFD+L-NAME comparison in Figure 1, at matched sample size of four to five mice per arm. (A) Spearman correlation between per-gene TAC-versus-sham effects in the two laboratories, across all shared genes and restricted to genes called differentially expressed in the GSE32277S arm, together with the eight-pathway summary; the corresponding HFD+L-NAME values from Figure 1 are plotted alongside for comparison. Error bars, bootstrap S5% confidence intervals. (B) Per-gene effects in the two TAC cohorts across the 1C,538 shared genes; shading, gene density; line, linear fit. (C) Genes reaching adjusted p below 0.05 in each reference cohort with four mice per group. (D) The same comparison summarized as eight pathway activity scores; circles, GSE32277S; triangles, GSE283181.

Sample size is not the binding constraint. With four mice per group, differential expression in the GSE322779 arm identified 2,241 genes at adjusted p below 0.05, against 349 in the internal HFD+L-NAME cohort at the same group size (Figure 2C). The framework therefore resolves gene-level concordance at this scale whenever the underlying effect is strong and reproducible.

The pathway level behaved differently. Summarizing the transverse aortic constriction comparison as the same eight pathway scores gave rho = 0.905, with all eight programs agreeing in direction (Figure 2D), against rho = 0.714 with six of eight agreeing for HFD+L-NAME. Both estimates are large, and the bootstrap interval for the transverse aortic constriction estimate spans 0.32 to 1.00 for the same reason as before: eight points constrain a correlation weakly. Pathway-level concordance was thus high both for a model that reproduces at gene level and for one that does not, whereas gene-level concordance separated the two by a factor of seven. This is the practical content of the resolution effect: pathway agreement locates shared biology, and gene-level agreement is what establishes that a model reproduces.

The pattern holds beyond these two models. I repeated the measurement on three further mouse cardiac models, each represented by two independent deposits carrying depositor-assigned labels: doxorubicin cardiotoxicity (GSE224157 and GSE307421), isoproterenol infusion (GSE217755 and GSE255532) and myocardial infarction (GSE275835 and GSE304090). Gene-level concordance ranged nearly nine-fold across the five model pairs, from rho = 0.078 for HFD+L-NAME and 0.153 for doxorubicin to 0.525 for isoproterenol, 0.535 for transverse aortic constriction and 0.687 for infarction. Pathway-level concordance did not track that range at all: every pair returned 0.714 or above, spanning only 0.714 to 1.000 (Figure S2A, B). The two doxorubicin cohorts make the point concretely. Both show the expected p53 and atrophy response, so both captured the model, yet they agreed at rho = 0.153 gene by gene while returning exactly the same pathway-level correlation, 0.714, as the HFD+L-NAME comparison.

### Cross-species concordance is confined to oxidative phosphorylation

I next compared per-gene effects in each murine model against subgroup-resolved human HFpEF myocardial proteomics. The pre-specified support filter prioritized three pathways with at least 15 shared genes: oxidative phosphorylation (142), collagen fibril organization (17) and mechanotransduction (17) (Figure 3). Comparisons resting on fewer than 15 shared genes remain displayed for completeness but were excluded from inference, creating a transparent evidence map for translational interpretation.

**Figure 3.**
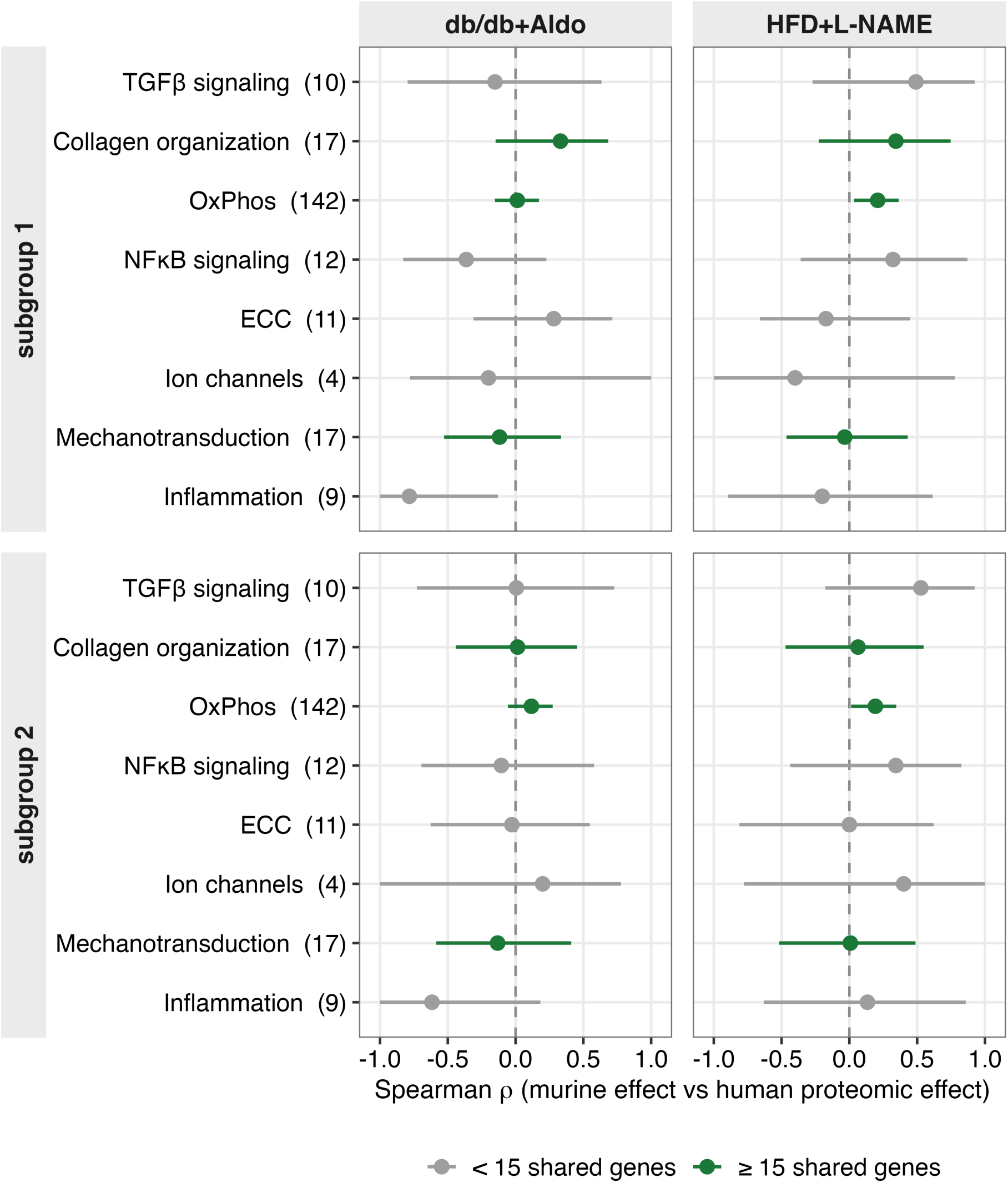
Murine-to-human pathway concordance and the gene support behind each estimate. Spearman correlation between per-gene murine HFpEF effects and human HFpEF myocardial proteomic effects, for both murine models against two human proteomic subgroups. Points are point estimates, bars bootstrap S5% confidence intervals, and the number of shared genes is given in parentheses. Comparisons resting on fewer than 15 shared genes (gray) were fixed in advance as uninterpretable and are shown only for completeness.

Within this supported set, oxidative phosphorylation was the only pathway to reach significance, doing so in the HFD+L-NAME model against both human subgroups (rho = 0.209, permutation p = 0.015 for subgroup 1; rho = 0.192, p = 0.023 for subgroup 2), with unadjusted bootstrap confidence intervals excluding zero. These nominal signals arise from a 12-comparison family and are hypothesis-generating; their recurrence across both human subgroups nevertheless makes oxidative metabolism the strongest cross-species candidate here, and separates HFD+L-NAME from db/db+Aldo for this specific biology.

### Bulk myofibroblast signatures track mural cells rather than fibroblasts

Bulk marker-based enrichment is widely used to attribute transcriptomic change to particular cell populations. I examined whether the curated signatures used for this purpose can support that attribution.

The independent single-cell projection provided a direct specificity test for the six-gene myofibroblast set (ACTA2, LOX, POSTN, TAGLN, THBS1 and TIMP1). Its composition combines canonical smooth-muscle markers (ACTA2 and TAGLN), a gene shared with the fibroblast set (POSTN), and broadly expressed matrix genes (LOX, THBS1 and TIMP1), without activated-fibroblast markers such as CTHRC1, COMP or FAP. Consistent with that composition, the signature scored highest in smooth muscle and pericytes (0.541) rather than annotated fibroblasts (0.184; Figure 4A). Composition and projection together show that this score is best interpreted as a mural/matrix-remodeling program rather than a fibroblast-specific state, converting an ambiguous bulk label into a more biologically precise readout.

**Figure 4.**
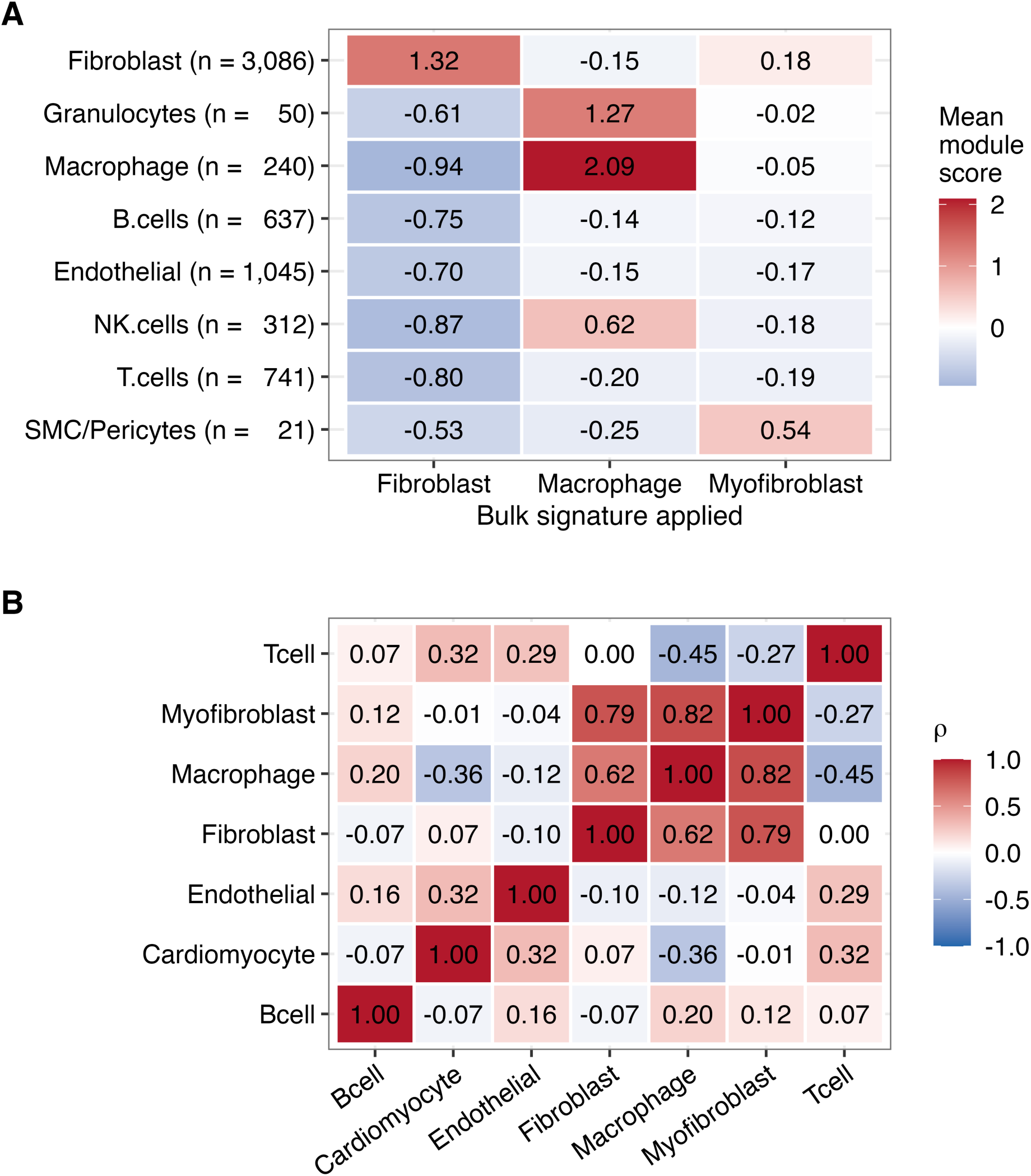
Resolution limits of bulk cell-program inference. (A) Mean module score for each curated bulk signature within each annotated cell type of the independent ventricular single-cell dataset GSE275031; cell numbers in parentheses. The myofibroblast signature scores highest in smooth muscle and pericytes rather than in fibroblasts. (B) Spearman correlation among bulk cell-program scores across the 15 db/db+Aldo samples, showing that the three fibro-inflammatory programs are not independent of one another.

The fibroblast and macrophage signatures behaved as intended, scoring highest in their named populations. Their scores were nonetheless strongly intercorrelated across samples (fibroblast-myofibroblast rho = 0.79, myofibroblast-macrophage rho = 0.82, fibroblast-macrophage rho = 0.62; Figure 4B), so they should be read as facets of a single fibro-inflammatory axis rather than as independent lines of evidence. Pathway scores shared substantial variance with the same axis (rho = 0.37 to 0.66). Transcription-factor activities were less reducible to it: only 23% of 273 inferred activities exceeded a squared rank correlation of 0.25 against the fibro-inflammatory composition axis (median 0.097), indicating that the regulatory layer carries information beyond bulk composition even though the pathway and cell-program layers largely do not.

Two of seven cell programs were significantly enriched in HFpEF after Benjamini-Hochberg correction: macrophage (adjusted p = 0.041) and myofibroblast (adjusted p = 0.049), with the fibroblast program showing a non-significant trend (adjusted p = 0.168). Read together with the specificity analysis above, the macrophage program is the one enriched signal that both survives correction and demonstrably identifies its named population.

### Cell programs and transcription factors, but not pathways, predict remodeling severity

Nested leave-one-out validation showed positive out-of-sample predictive skill for models using cell programs, transcription-factor activities or combined features (1 − SSE/SST = 0.31–0.43), whereas pathways alone performed below the sample mean (−0.19; Figure S1A). Squared correlation obscured this failure mode (Figure S1B). Matching the transcription-factor candidate pool to the eight pathway features removed the apparent transcription-factor advantage (median root-mean-square error, RMSE, 0.98; Figure S1C), demonstrating why candidate-pool size must be considered when comparing feature classes.

Which features the model selects is itself explained by cell composition. Across 233 transcription factors, bootstrap selection frequency correlated with dependence on the fibro-inflammatory composition axis (Spearman rho = +0.62, p = 3 × 10⁻²⁶; Figure S1D). The six factors retained in at least half of 500 refits had a median squared rank correlation of 0.55 against that single axis, against 0.12 for the remainder (Wilcoxon p = 0.0001), and five of them rank among the fifteen most composition-dependent of 273. Regularized selection at this sample size therefore recovers whichever features track bulk cell content rather than regulators with independent explanatory value, which is the expected behavior when one axis dominates the sample-level variance and accounts for the absence of a cardiac rationale among the top-ranked factors.

### Results are robust to sex and to body-size normalization

The cohort is close to sex-balanced (4 female and 4 male HFpEF, 4 female and 3 male control; Fisher exact p = 1.000), no pathway showed a sex effect at p below 0.05, and group differences are therefore not attributable to sex imbalance. Because db/db animals are markedly obese, echocardiographic left ventricular mass could track body size rather than disease; it correlates with body weight at rho = 0.71. Repeating the analyses with heart weight normalized to tibia length, the standard body-size-independent index available in the source morphometry, gave a severity score correlating with the original at rho = 0.95 and left every cell-program association essentially unchanged (Figure S3A). Partialling out body weight did not weaken the associations. Benjamini-Hochberg correction across the seven cell programs retained macrophage and myofibroblast enrichment, while the remaining programs did not meet the adjusted threshold (Figure S3B). No pathway-level group difference survived correction across the eight pathways.

## Discussion

This study establishes resolution-aware benchmarking as a practical strategy for extracting reproducible and translatable signals from heterogeneous HFpEF omics datasets. By integrating gene-, pathway- and cell-program evidence, the framework locates where concordance is strongest and converts apparently divergent results into a coherent hierarchy of biological evidence.

Across independent laboratories, the strongest transcriptional effects showed substantial agreement and nearly three quarters reproduced in direction. Pathway summarization returned a coordinated signal across six of eight programs, on an estimate whose interval spans −0.04 to 0.98 because eight points constrain a correlation weakly. The two resolutions answer different questions: gene-level results define the reproducible transcriptional core, and pathway-level results organize that core into conserved and context-sensitive programs.

Inconsistency between transcriptomic studies of the same disease is well documented: a consensus analysis of sixteen human end-stage heart failure cohorts found that individually reported genes agree poorly between studies even where a meta-analytic signature is recoverable [26]. The present results locate where that inconsistency resides. Claims of cross-dataset reproducibility should include gene-level agreement, which provides a denser description of the effect distribution, while recognizing that correlated genes are not independent biological replicates. Pathway-level summaries remain useful for interpretation but should not serve as the sole evidential basis. Where pathway-level correlations are reported, the number of contributing features and an interval estimate should accompany them as a matter of course.

A positive control makes the resolution effect actionable. Two independent transverse aortic constriction cohorts, analyzed at the same sample size with the same pipeline, agreed at rho = 0.54 across shared genes, whereas the HFD+L-NAME comparison agreed at rho = 0.08, and four mice per group were sufficient to call more than two thousand differentially expressed genes. Pathway-level correlations, by contrast, were high in both comparisons. The discriminating measurement is therefore gene-level agreement, and a pathway correlation on its own does not distinguish a model that reproduces across laboratories from one that does not. This supplies a concrete benchmark for model-selection studies: report gene-level concordance alongside a positive control profiled at comparable size. The same measurement across three further model pairs, doxorubicin, isoproterenol and myocardial infarction, returned the same picture: gene-level agreement ranged from 0.15 to 0.69, while every pair returned a pathway-level correlation of 0.71 or above (Figure S2).

Cross-species benchmarking yielded one well-supported result. Oxidative phosphorylation in the HFD+L-NAME model correlated with both human HFpEF subgroups, and the feature-support filter established which further pathways could be evaluated at all. This shifts model selection from asking whether a model is generically human-like to identifying the model best matched to a defined biological question.

Notably, sample group labels are assumed correct by every downstream statistic, and an error in them is invisible to any amount of subsequent statistical care. Public deposits of the kind used here almost always carry the means to check: an accompanying phenotype table, or the depositors’ own differential expression results. Verifying labels against at least one such independent source costs little and should be routine in re-analysis work; I have made it a required step in the released pipeline.

Independent single-cell validation adds biological resolution to the bulk findings. The myofibroblast score is best interpreted as a mural/matrix-remodeling program, whereas the fibroblast and macrophage signatures retain their intended specificity. Human single-nucleus data reinforce the value of this distinction: in septal myocardial biopsies from 19 HFpEF patients and 24 non-failing controls, fibroblasts carried more differentially expressed genes than any other cell type, while the transcriptional features most specific to HFpEF were found in cardiomyocytes [27]. Together, these results support a cell-state-aware view of HFpEF remodeling: fibro-inflammatory activity is prominent, macrophage enrichment is robust after correction, and bulk signatures become more biologically informative when anchored to independent single-cell data.

This study advances both methodology and translation. It integrates cross-laboratory replication, cross-species feature support, uncertainty calibration and independent single-cell validation into one reproducible workflow. The result is a generalizable framework for converting public omics into decision-ready evidence for disease-model selection, and its HFpEF application delimits a reproducible gene-level core, one supported cross-species pathway, and the cell programs that bulk signatures can and cannot resolve.

Several constraints bound these conclusions. The cross-laboratory comparisons rest on four to six animals per arm, at or below the replicate numbers recommended for bulk RNA-sequencing differential expression [28]; the positive control establishes that this is sufficient to resolve a strong and reproducible effect, while leaving smaller effects beyond reach. Cell states were inferred from bulk tissue and tested against independent single-cell data rather than measured directly. Feature-resampling intervals describe stability to gene and pathway composition and do not substitute for replication in new animals. The eight-point pathway estimator is weakly determined by construction, which is the property measured here rather than a defect of any one dataset. The cross-species comparison relates murine transcript effects to human protein effects, so species and molecular layer cannot be separated in these estimates.

The study design also defines clear next applications. Additional cohorts can refine the numerical estimates and extend the framework to other HFpEF models, while matched transcript–protein datasets can separate species effects from molecular-layer effects. Feature-resampling intervals quantify stability to gene and pathway composition; independent animal and cohort replication remains the complementary next level of validation. The supplementary severity models are exploratory because the cohort is modest, but nested cross-validation and candidate-pool matching provide a rigorous proof of concept. For cell-program specificity, marker composition and independent single-cell projection provide convergent evidence despite the small mural population.

For the reduction and replacement of animal use, the framework offers an immediately deployable form of public-data triage. By identifying reproducible effects and human-aligned pathways before experiments are commissioned, it concentrates animal studies on the best-supported biological questions and reduces duplicative or weakly motivated work. Between-laboratory variation is itself a recognized driver of poor reproducibility in animal research[29], and measuring it directly in public data is a low-cost way to anticipate it. Because the inputs are public and the pipeline is reproducible, the approach can be applied broadly across preclinical disease-model programs.

## Data and code availability

Murine RNA-sequencing and phenotype data were obtained from the Gene Expression Omnibus (GEO) under GSE284354; external validation RNA-sequencing from GSE180065; single-cell data from GSE275031; positive-control transverse aortic constriction data from GSE322779 and GSE283181; cross-model generality cohorts from GSE224157, GSE307421, GSE217755, GSE255532, GSE275835 and GSE304090; and human HFpEF proteomics from the public deposit described in [8]. All analysis code, curated pathway gene sets, cell-type marker lists and the transcription-factor regulon table will be released at https://github.com/aminforouzandehmehr and archived at Zenodo upon publication, and are sufficient to reproduce every figure and table from the public source files. They are available to editors and reviewers on request during peer review. No new data were generated for this study.

## Author contributions

A.F. conceived the study, designed and performed all analyses, prepared the figures, and wrote the manuscript.

---

**Figure S1.**
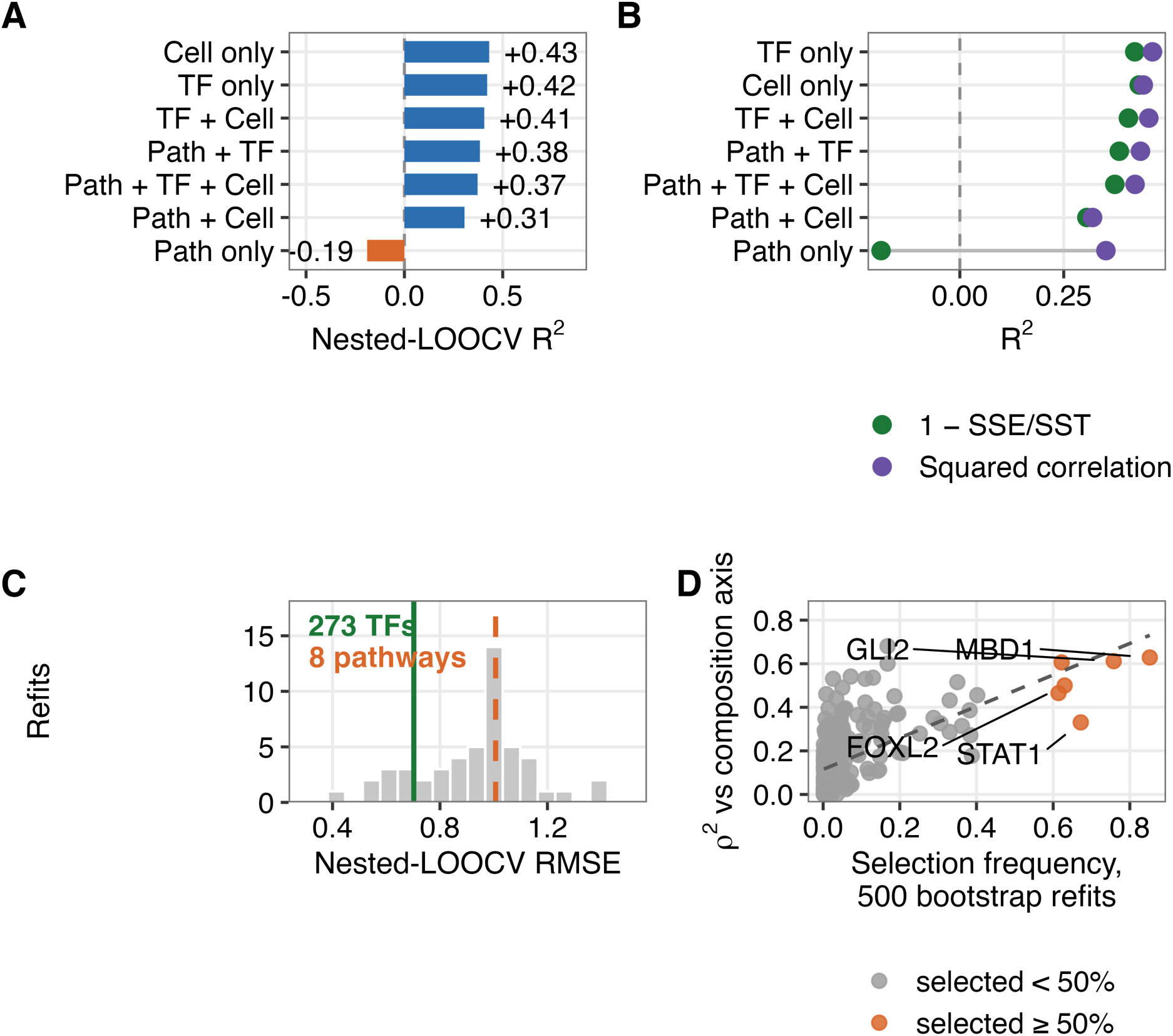
Phenotype-derived severity modeling in 15 mice. (A) Out-of-sample predictive skill by feature class, computed as 1 − SSE/SST; values below zero indicate performance worse than predicting the sample mean. (B) The same fits scored by 1 − SSE/SST and by the squared observed-predicted correlation, which cannot fall below zero and therefore cannot reveal a model without predictive skill. (C) Distribution of nested leave-one-out cross-validated RMSE across 50 refits using eight randomly chosen transcription factors, matching the number of pathway features, against the full 273-feature transcription-factor model and the pathway model. (D) Bootstrap selection frequency versus dependence on the fibro-inflammatory composition axis for 233 transcription factors; the most frequently selected factors are among the most composition-dependent. Feature classes are abbreviated in panels A and B: Path, pathways; TF, transcription factors; Cell, cell programs.

**Figure S2.**
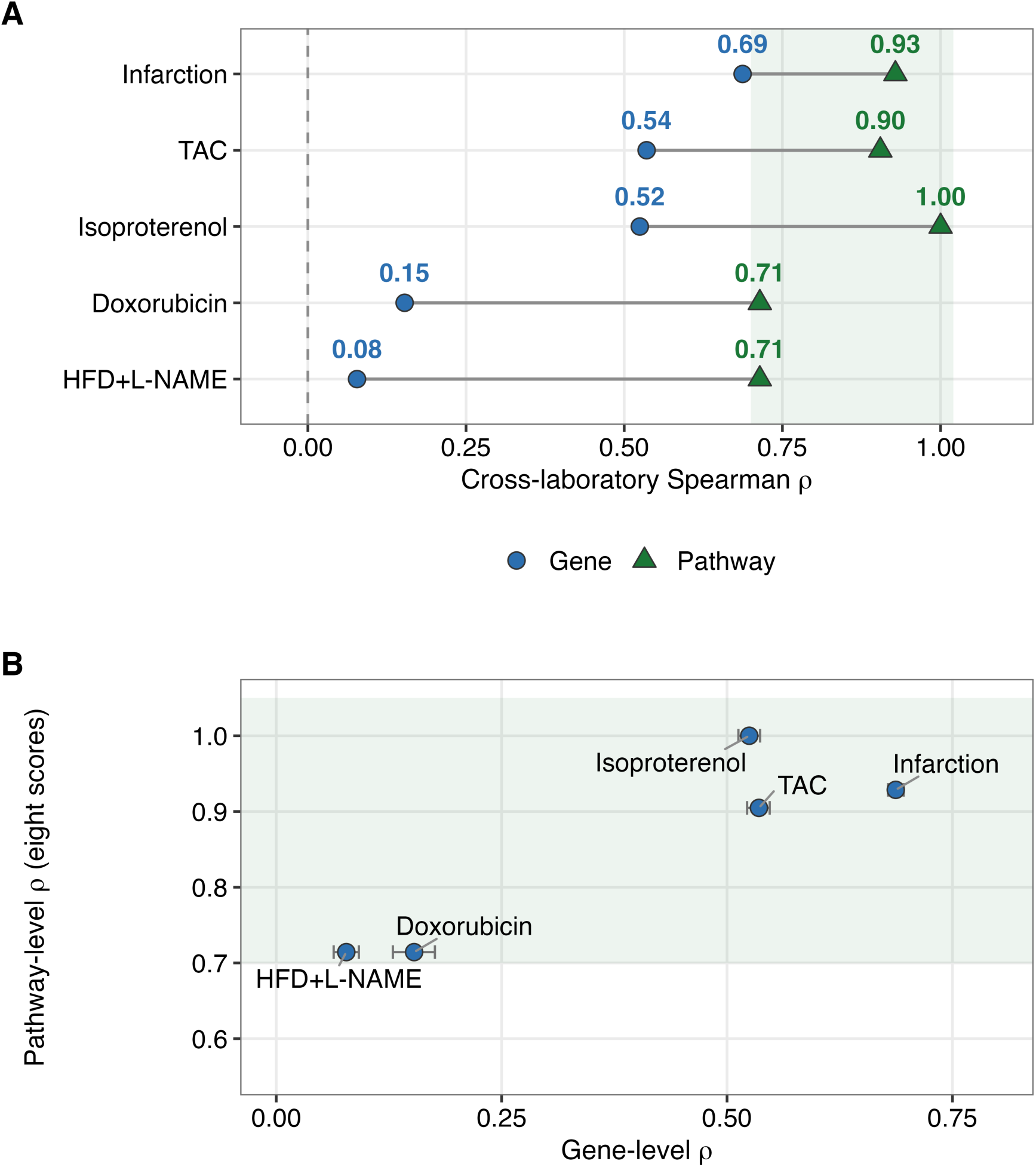
The resolution effect across five mouse cardiac models. Each model was measured in two independent deposits using the pipeline applied in Figures 1 and 2: doxorubicin (GSE224157 and GSE307421), isoproterenol (GSE217755 and GSE255532) and myocardial infarction (GSE275835 and GSE304090), together with the transverse aortic constriction and HFD+L-NAME pairs from the main text. All group labels are the depositors’ own. (A) Cross-laboratory Spearman correlation at gene level (circles) and for the eight-pathway summary (triangles), ordered by gene-level agreement; shading marks pathway correlations above 0.70. (B) The same five pairs plotted against one another; horizontal bars are bootstrap 95% confidence intervals on the gene-level estimate; shading as in A.

**Figure S3.**
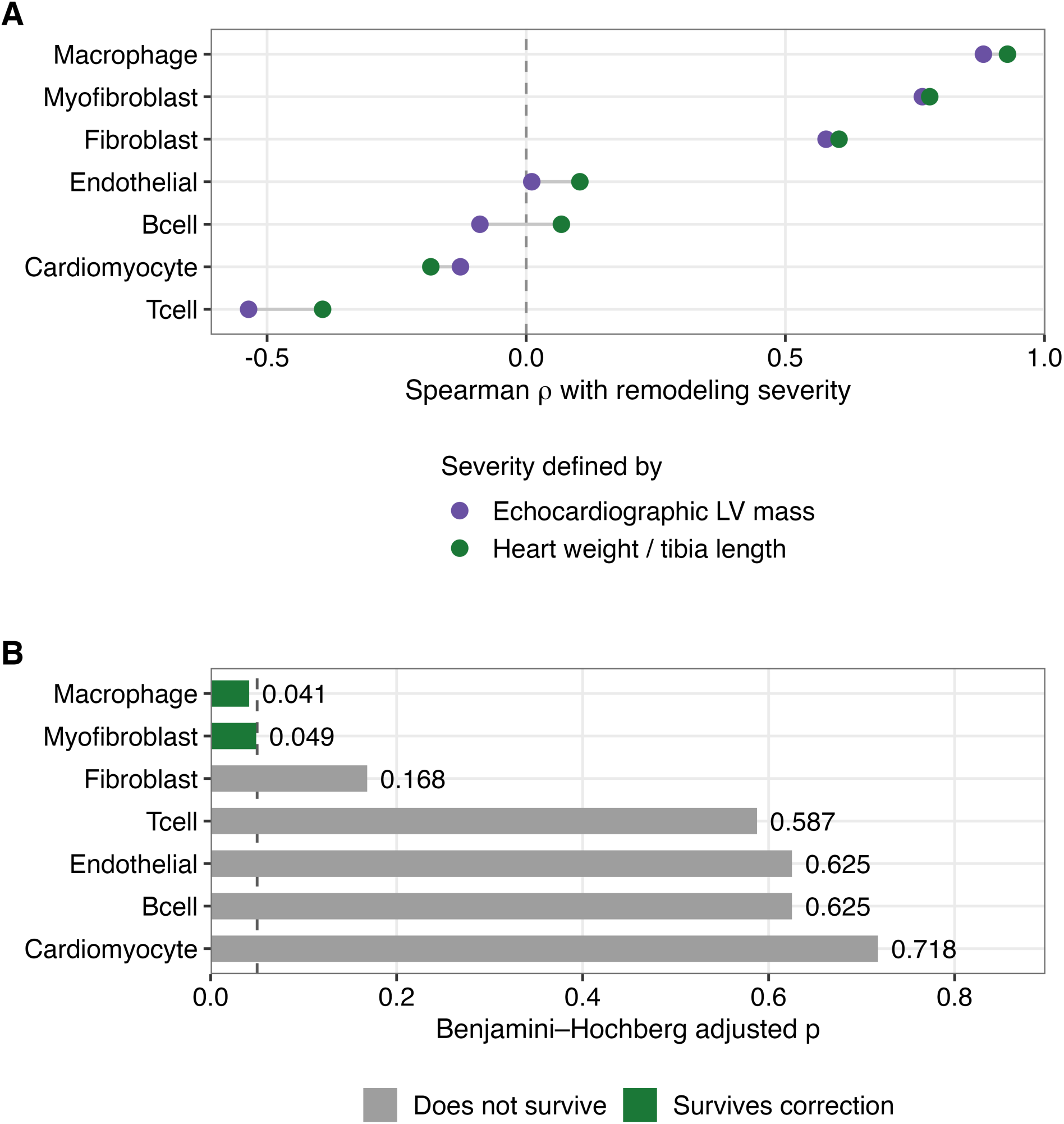
Sensitivity analyses. (A) Association between each cell-program score and remodeling severity, with severity defined from echocardiographic left ventricular mass and, alternatively, from heart weight normalized to tibia length. (B) Benjamini–Hochberg adjusted p-values for cell-program enrichment in HFpEF versus control (Wilcoxon rank-sum test), corrected across all seven programs.

